# Relationships between GABA+ and Glx concentrations with age and inhibition in healthy older adults

**DOI:** 10.1101/2025.02.25.640218

**Authors:** Ciara Treacy, Sophie C. Andrews, Jacob M. Levenstein

## Abstract

Inhibition represents a core executive function which underlies the ability to suppress interfering and/or distracting stimuli, thereby building resistance against task-irrelevant information. However, the impact of ageing on inhibitory functioning and the potential role of neuroplasticity - largely driven by predominant excitatory (glutamatergic) and inhibitory (GABAergic) neurochemicals – remains poorly understood. In this study, we investigated age relationships with neurochemical concentrations (GABA+ and Glx) and examined the associations between these neurochemicals and inhibitory sub-components in the context of healthy ageing. To this end, participants completed three inhibition tasks (i.e., flanker, Stroop and go/no-go), each measuring a different sub-component process, via the PsyToolkit platform. MRS data was acquired in the sensorimotor (SM1; n=71, mean age (SD) = 68.3 (±9.7) years, 39 females) and prefrontal (PFC; n=58, mean age (SD) = 67.6 (±9.6) years, 30 females) regions using a HERMES sequence and analysed using OSPREY’s pipeline. After correcting for gender and education, semi-partial correlations (*rho*) revealed no significant relationships between age and GABA+ or Glx concentrations in either the SM1 or PFC regions. Furthermore, through partial correlations (*rho*), after correcting for age, gender and education, we identified a significant negative relationship between SM1 Glx concentrations and go/no-go error rates, such that greater concentrations of Glx in the SM1 region were associated with greater accuracy on the go/no-go task. The null age-neurochemical results suggest that GABA+ and Glx may not uniformly decline during healthy ageing. This finding suggests that the relationship between older age and neurochemistry may be more nuanced than previously reported. In addition, our neurochemical-behavioural findings provide neurochemically-and-spatially specific evidence that SM1 Glx concentrations may be important for response inhibition. This result indicates a role for the glutamatergic system in supporting inhibition over the normal course of ageing.

## 1.0 Introduction

Successful inhibitory functioning relies upon the ability to voluntarily suppress impulses toward environmental interference and ignore competing distractions, thereby facilitating goal-oriented behaviour (Diamond, 2013). Debate persists regarding the impact of ageing on inhibitory functioning, with behavioural evidence supporting an age-related decline (Andrés et al., 2008; Colcombe et al., 2005; Fong et al., 2021; Kouwenhoven & Machado, 2024; Nielson et al., 2002; Zhu et al., 2010), and refuting an ageing influence (Borella et al., 2009; Fernandez-Duque & Black, 2006; Grandjean & Collette, 2011; Hsieh & Fang, 2012; Salthouse, 2010). Two independent meta-analytical reviews exemplify these contradictory findings and challenge the hypothesis of a general inhibition deficit in ageing, with Verhaegen (2011) failing to identify any specific age-related deficits in inhibition across eight tasks, whilst Rey-Mermet and Gade (2018) detected age-related decrements in only a select number of the eleven evaluated inhibitory tasks. Notably, these meta-analyses identified at least eight different tasks to measure inhibition, highlighting the diverse range of task designs and stimuli employed to evaluate age-related changes. This variability in methodology may be obscuring relationships associated with healthy ageing, exacerbating the inconsistencies observed in the literature.

Cognitive changes observed during normal ageing can, in-part, be explained by neuroplasticity (Burke & Barnes, 2006), a process largely driven by the relative balance of predominant excitatory and inhibitory neurochemicals (Duncan et al., 2014). The ratio of excitatory and inhibitory neurons within cortical circuits is under precise, homeostatic control to generate a stable global level of activity (Sohal & Rubenstein, 2019). Within the cortex, glutamate (Glu) and γ-aminobutyric acid (GABA), the major excitatory and inhibitory neurochemicals, respectively, are closely related to the excitation-inhibition (E/I) balance (Isaacson & Scanziani, 2011). GABA is largely responsible for cortical network synchrony, neurogenesis and preventing neural network hyperactivity by diminishing the prospect of a post-synaptic neuronal firing (Jiménez-Balado & Eich, 2021; Xu et al., 2020). Glu, a neurotransmitter and substrate of GABA synthesis, functions as the primary mediator of excitatory signals, neuronal plasticity, and synaptogenesis by depolarising post-synaptic membranes, which increases the likelihood of an action potential firing (Gao & Penzes, 2015). Magnetic resonance spectroscopy (MRS) is a non-invasive in-vivo imaging technique to quantify neurochemical concentrations in specific brain regions, with edited experiments capable of isolating low-concentration neurochemicals with substantial signal overlap, such as GABA (often denoted as “GABA+” due to co-editing of macromolecules at similar resonances; see Saleh et al. (2019)).

MRS quantification of Glu presents a similar challenge due to the significant signal overlap between Glu and glutamine (precursor molecule) resonances, thus, their contributions are often combined and referred to as “Glx” (Mullins et al., 2014; Ramadan et al., 2013). Disruptions to the neural E/I balance appears to play a detrimental role in the pathogenesis of neurodegenerative diseases of ageing, such as Alzheimer’s Disease (Bi et al., 2020). In healthy adults, E/I (im)balances have been associated with working memory (Marsman et al., 2017), conflict resolution (de la Vega et al., 2014), perceptual performance (Kondo et al., 2018) and inhibitory functioning (Koizumi et al., 2018). Consequently, the crucial roles of GABA+ and Glx in coordinating neural function and stability (Bannai et al., 2020) has prompted MRS investigations to examine their presentation in the ageing brain and role in human behaviour.

While research investigating the impact of healthy ageing on MRS-assessed neurochemical concentrations remains limited, older adults typically exhibit reduced GABA+ and Glu (and/or Glx) concentrations across the brain (for meta-analytic reviews, see Porges et al. (2021) and Roalf et al. (2020)). For example, age-related declines in cortical neurochemical concentrations have been reported across the cerebral cortex for GABA+ (Gao et al., 2013; Porges, Woods, Edden, et al., 2017; Simmonite et al., 2019) and Glx (Ding et al., 2016), including the motor regions (GABA+, Grachev and Apkarian (2001); Hermans et al. (2018); Zuppichini et al. (2024); Glu, Kaiser et al. (2005); Liu et al. (2024). Furthermore, the relationship between MRS-assessed GABA and Glu (and/or Glx) concentrations and behavioural inhibition in healthy older adults (i.e., ≥ 50 years) remains unclear due to limited evidence. Of these few studies includes work conducted by Hermans and colleagues (2018) who identified that greater SMA GABA+ concentrations were associated with greater reactive inhibition, as measured using a stop signal task (SST). Furthermore, recent work by Liu et al. (2024) examined 18 younger (age range = 20-35 years; M_age_ = 22.3 (±1.3) years) and 17 older (age range = 65-85 years; M_age_ = 71.1 (±2.6) years) healthy adults and reported that Glu concentrations negatively correlated with reactive inhibition performance (as measured using the SST) in the younger group, however, this relationship was not observed in the older group. Drawing on additional MRS investigations of inhibition in younger adults, GABA+ concentrations have been positively associated with response inhibition (as measured using the go/no-go task) in the striatum (Quetscher et al., 2015), anterior cingulate cortex (Silveri et al., 2013) and superior temporal gyrus (Cheng et al., 2017). Beyond inhibition, GABA and Glu (and/or Glx) concentrations have been cross-sectionally related with other cognitive abilities in healthy older adults. For example, neurochemical concentrations have been positively associated with motor performance (GABA+ (from SM1), Cassady et al. (2019)), sensory discrimination (GABA+(from SM1), Puts et al. (2011)), general cognitive function (GABA+ (from frontal and posterior; superior to the splenium and aligned with corpus callosum), Porges, Woods, Edden, et al. (2017)), fluid processing (Glx (from visual cortex), Simmonite et al. (2019)), working memory and decision making (Glu (from prefrontal), Rmus et al. (2023)). So far, preserving GABA+ and Glu (and/or Glx) concentrations in the ageing brain appears to invariably relate to better cognitive outcomes, however further work is needed to confirm these associations across brain regions and tasks in a single study population of healthy older adults. Elucidating the role of GABA+, Glx and their relative (im)balance in inhibitory functioning may help explain the behavioural variability prevailing in ageing.

Prior MRS investigations of inhibition primarily use a singular task to measure response inhibition (e.g., go/no-go or SST), which limits comprehensive assessment of inhibitory functioning, and may obscure behavioural relationships with GABA+ and Glx concentrations. An often-overlooked consideration is that behavioural inhibition is not unitary, for example, Rey-Mermet and Gade (2018) propose three sub-components of inhibition: the ability to inhibit i) distracting information (i.e., flanker), ii) response interference (i.e., Stroop), and ii) prepotent responding (i.e., go/no-go). In healthy older adults, these inhibitory sub-components appear to be distinct from one-another and show variable age effects (Treacy et al., under-review), therefore, it is probable that they are subserved by distinct mechanisms with different neurochemical dependencies. Additionally, in MRS studies important a priori assumptions are made regarding voxel of interest (VOI). Insight from fMRI studies identify the fronto-parietal network as the core system engaged in inhibition, with clusters in the frontal cortex, angular gyrus and SMA (for a meta-analytic review, see Zhang et al. (2017)). Furthermore, inhibiting a prepotent (motor) response also depends upon the fronto-basal ganglia network, including the SM1 (Aron, 2011). While prior studies have linked SM1 and PFC GABA+/Glx concentrations to various cognitive domains (for reviews see Li et al. (2022); Roalf et al. (2020)), neurochemical links to inhibitory sub-components have not been investigated in healthy older adults specifically (≥ 50 years).

To address this gap, we clarify the relationships between SM1 and PFC neurochemical concentrations and three sub-components of behavioural inhibition in healthy older adults aged ≥ 50 years. For aim 1, we examined the relationship between healthy ageing, GABA+ and Glx concentrations from the SM1 and PFC. We hypothesised that age would be negatively associated with GABA+ and Glx concentrations in both brain regions, such that older age would relate to lower neurochemical concentrations. For aim 2, we investigated relationships between SM1 and PFC neurochemical concentrations and behavioural inhibition, as measured using the flanker, Stroop and go/no-go tasks. We hypothesised that GABA+ and Glx concentrations would be negatively associated with performance on all three tasks in both brain regions, such that higher neurochemical concentrations would relate to greater inhibitory abilities. As an exploratory aim, we assessed these ageing and behavioural relationships using the E/I ratio. Specifically, we hypothesised a positive relationship between ageing and the E/I ratio, such that older age would relate to a greater E/I imbalance in both brain regions. To clarify, although we hypothesise both lower GABA+ and lower Glx concentrations during ageing, the rate of age-related decline for these neurochemicals is not well understood and may not decline at the same rate (Thomson et al., 2024). Lastly, we hypothesised that the E/I ratio would be positively associated with performance on all behavioural tasks, such that a greater E/I imbalance would relate to worse inhibitory abilities.

## 2.0 Methods

### 2.1 Participants

This study was approved by the Human Research Ethics Committee of the University of the Sunshine Coast (S211620), with data collection occurring at the UniSC Thompson Institute. All participants provided written, informed consent. Healthy older adults aged 50-85 years were recruited from the general community through a study specific webpage, media announcements and local community groups. Eligible participants were right hand dominant, English-speaking, generally healthy individuals absent of any diagnoses pertaining to mild cognitive impairment, psychiatric disorders (e.g., schizophrenia or bipolar), or major neurological conditions (e.g., dementia or Parkinson’s). Additionally, participants were excluded if they presented with MRI contraindications (e.g., pacemaker, stents, metallic foreign bodies or weight exceeding 120kg), colour-blindness, respiratory conditions (e.g., COPD), cardiovascular conditions (e.g., stroke, TIA or myocardial infarction), prior head injuries (loss of consciousness >60mins), poorly controlled diabetes, current substance abuse or misuse, or usage of medications known to impact the central nervous system (e.g., antidepressant or antianxiety medication). Eligibility criteria were operationalized via online self-report questionnaires and telephone screening. In total, 81 individuals satisfied eligibility screening and were enrolled in the study. Of these, two individuals did not complete the MRI due to scanning contraindications, two individuals had incomplete spectroscopy datasets (n=1 incomplete SM1, n=1 incomplete PFC), and one individual was excluded post-hoc due to unreported cognitive difficulties impacting task compliance. Therefore, the MRS cohort consisted of 77 participants. After data quality and accuracy verification, the final MRS sample was reduced to 71 participants for the SM1 dataset (M_age_ = 68.2 years, 39f) and 58 participants for the PFC dataset (M_age_ = 67.6 years, 30f), see Table 1 for univariate statistics.

**Table 1:** Participant sample characteristics. n (sample size); female(f); male(m); sensorimotor (SM1); prefrontal cortex (PFC); relative to total creatine (/tCr); excitation/inhibition (E/I); balanced integrated score (*bis*).

|  | **n** | **Mean(±SD)** | **Median** | **Min - Max** |
| --- | --- | --- | --- | --- |
| ***SM1 Sample Characteristics*** |  |  |  |  |
| Age (years) | 71 | 68.28(±9.67) | 70.38 | 50.56 – 84.77 |
| Gender (f, m) | 39, 32 | - | - | - |
| Education (years) | 71 | 16.36(±3.79) | 16.00 | 9.00– 26.00 |
| GABA+/tCr | 68 | 0.38(±0.17) | 0.37 | 0.04 – 0.80 |
| Glx/tCr | 71 | 1.02(±0.32) | 1.00 | 0.43 – 1.83 |
| E/I ratio | 68 | 3.65(±2.94) | 2.50 | 0.66 – 14.68 |
| Go/no-go (error rate) | 70 | 9.63(±8.54) | 6.67 | 0.00 – 34.33 |
| Go/no-go (bis) | 70 | 2.09(±1.15) | 2.11 | -3.47 – 2.09 |
| Stroop Effect (reaction time) | 71 | 88.95(±65.99) | 73.00 | -11.50 – 290.00 |
| Stroop Effect (error rate) | 71 | 1.48(±3.30) | 0 | -3.33 – 13.43 |
| Flanker Effect (reaction time) | 71 | 22.96(±30.84) | 28.00 | -73.00 – 109.00 |
| Flanker Effect (error rate) | 71 | 0.86(±2.86) | 1.28 | -7.37 – 8.00 |
| ***PFC Sample Characteristics*** |  |  |  |  |
| Age (years) | 58 | 67.60(±9.55) | 69.84 | 50.56 – 84.77 |
| Gender (f, m) | 30, 28 | - | - | - |
| Education (years) | 58 | 16.27(±3.63) | 16.00 | 9.00 – 26.00 |
| GABA+/tCr | 47 | 0.33(±0.16) | 0.34 | 0.04 – 0.68 |
| Glx/tCr | 57 | 1.32(±0.34) | 1.35 | 0.53 – 2.06 |
| E/I ratio | 46 | 5.83(±5.71) | 3.77 | 0.78 – 26.86 |
| Go/no-go (error rate) | 58 | 9.50(±8.11) | 6.67 | 0.00 – 33.33 |
| Go/no-go (bis) | 58 | 1.90(±1.02) | 2.00 | -1.98 – 1.90 |
| Stroop Effect (reaction time) | 58 | 82.86(±54.67) | 73.75 | -11.50 – 205.50 |
| Stroop Effect (error rate) | 58 | 1.61(±3.75) | 0 | -3.33 – 15.10 |
| Flanker Effect (reaction time) | 58 | 24.86(±31.31) | 32.50 | -73.00 – 109.00 |
| Flanker Effect (error rate) | 58 | 0.41(±2.72) | 0.85 | -6.77 – 5.41 |

### 2.2 Behavioural task paradigms

The specifics of the behavioural paradigms used in the present study have been reported elsewhere (Treacy et al., under-review). In short, a 5-minute letter flanker task (150 experimental trials, approximately equal proportion of congruent and incongruent trials), 4-minute Stroop colour and word task (120 experimental trials, equal proportion of congruent and incongruent trials), and 4-minute go/no-go task (225 experimental trials, frequent go events and rare no-go stimuli in a ratio of 4:1) were administered via the PsyToolkit platform v3.4.2 (Stoet, 2010, 2017), see Figure 1 for task diagrams. These tasks represent sub-component processes of inhibition, namely, the ability to inhibit distraction (flanker), interference (Stroop), and prepotent responding (Go/no-go). Upon arrival at the UniSC Thompson Institute, participants were seated in a quiet room at arm’s length from a DELL laptop screen (15.6” Precision 5530 model, intel UHD graphics 630), which was used to administer the PsyToolkit behavioural experiments. The aforementioned task timings reflect the “real-test” period, task instructions and practice trials are not included in these time estimations. Participants could take a short break (∼ 5 minutes) in between each task before the next set of instructions were given. For each task, participants were directed to maintain finger contact with the response key/s for the duration to ensure accurate record of reaction times and detection of slips of action. Further, participants were also instructed to give equal importance to both the speed and accuracy of their responses. These behavioural experiments were administered within 1-hour of MRI scanning.

**Figure 1:**
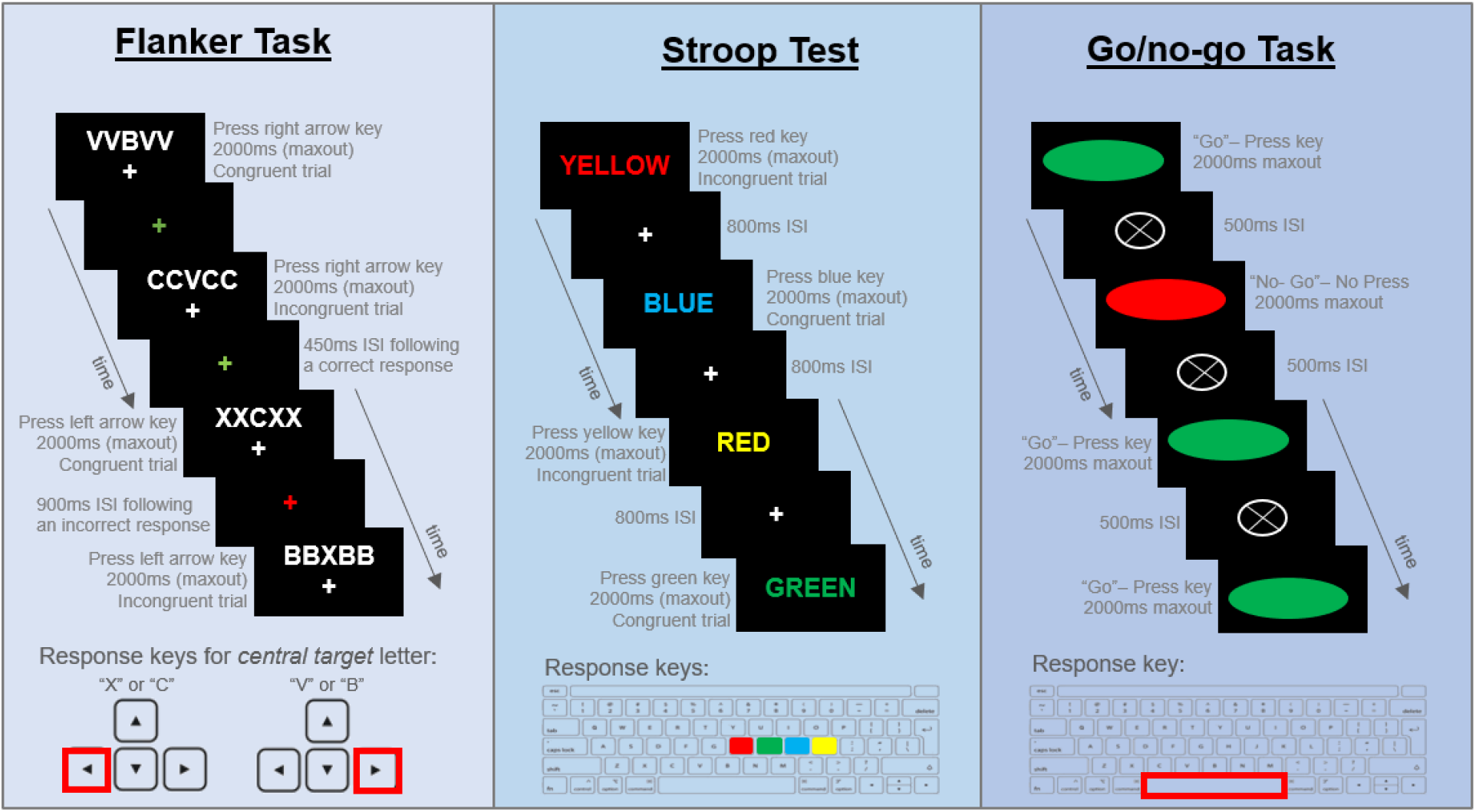
Behavioural task diagrams for the flanker, Stroop and go/no-go inhibitory measures, adapted from Treacy et al. (under-review). Inter-stimulus-interval (ISI).

We computed standard performance outcome measures related to inhibition and, where appropriate, speed-accuracy trade-off scores. Specifically, performance on the congruence tasks (i.e., flanker and Stroop) was measured by comparing congruent and incongruent trials in terms of median reaction times and error rates, calculating a congruence effect difference score for each. Go/no-go performance was measured using error rate (i.e., proportion of commission errors) and a balanced integrated score (bis; Liesefeld and Janczyk (2019)), which accounts for speed-accuracy trade-offs to maintain “real” effects. The bis was calculated using the standardised mean difference between the proportion of accurate responses and reaction times on correct trials (reaction times >700ms or < 200ms were removed), such that *bis* = z(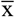 percentage correct) - z(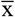 reaction time). As the bis was the only performance measure in this study where higher scores indicate better performance, this measure was reverse scored to facilitate hypothesis testing (i.e., higher scores on all behavioural measures represents poorer performance).

### 2.3 MRI data acquisition and processing

For all participants, MRI brain scans were performed on a 3T Siemens Skyra (Erlangen German) via a 64-channel head and neck coil. Following localisers, a T1-weighted magnetization-prepared rapid gradient echo (MPRAGE: TR =2200ms, TE=1.71ms, TI=850ms, flip angle=7°, voxel resolution=1mm^3^, FOV=208×256×256, PAT-GRAPPA=2, TA=3:57) was included to plan the MRS voxel placement. Single voxel MRS data were acquired in two different voxels of identical dimension using a Hadamard Encoding and Reconstruction of MEGA-Edited Spectroscopy sequence (HERMES) sequence (Chan et al., 2016), optimized for measuring GABA and GSH (TR=2000ms, TE=80ms, flip angle=90°, averages=320, edited pulse 1 = 1.9pmm, edited pulse 2 = 4.56ppm, edited off pulse = 7.5ppm, TA=10:48). Specifically, voxels were placed in the PFC and SM1 regions of the left cortex (Figure 2). The MRS post-processing and analysis procedures have been reported elsewhere, including specifics regarding data quality, model fit accuracy and case-wise/listwise spectra exclusions (Levenstein et al., under-review). Presented neurochemical concentrations are expressed as a ratio over creatine and phosphocreatine, combined as tCr, see Table 1 for descriptives.

**Figure 2:**
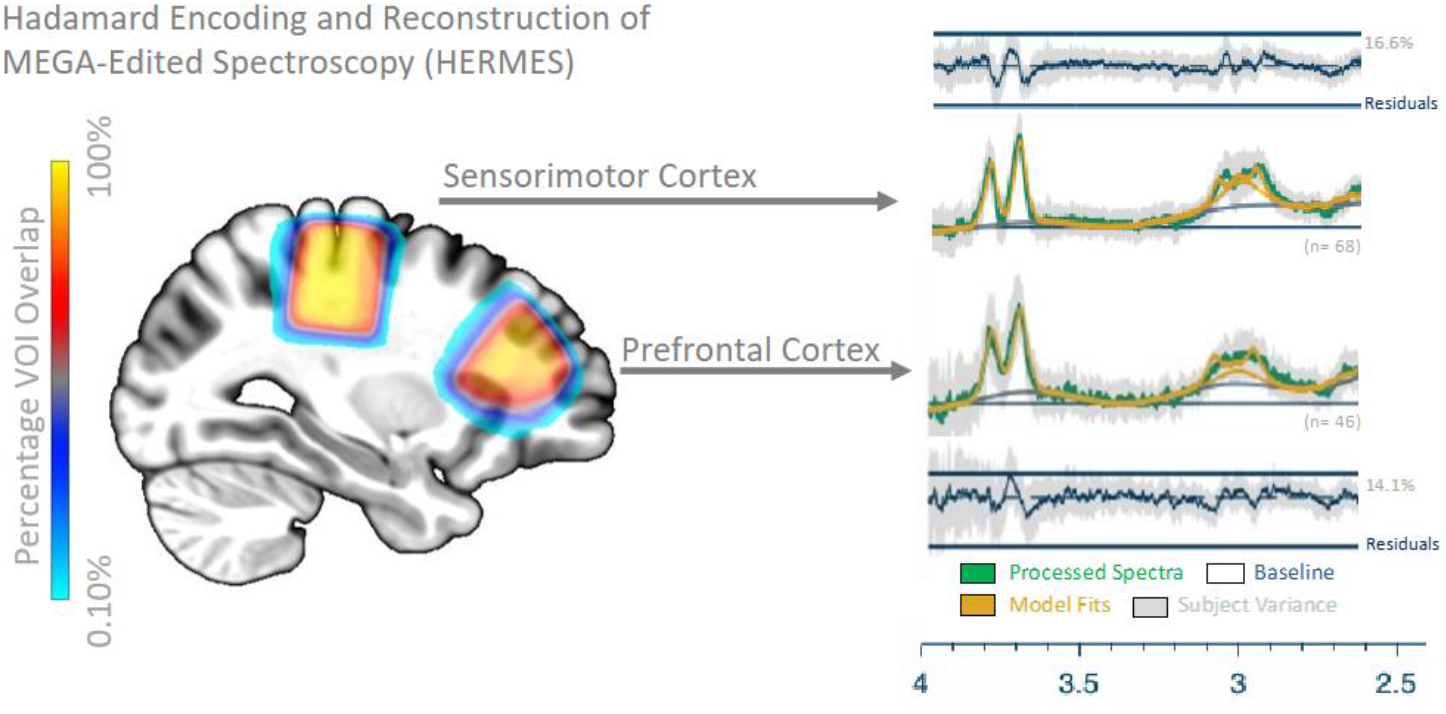
Frequency plot of MRS VOI locations in the left hemisphere (the SM1 and PFC) and corresponding model fits.

### 2.4 Statistical Analyses

All data statistical analyses were performed using R Statistical Software v4.3.2 (R Core Team, 2023). Outliers were identified using z-scores, with z-scores >3.29 or <3.29 considered as being extreme cases requiring correction. Outlier adjustment was performed as required using the z-score standard deviation transformation method, whereby extreme cases were adjusted to one unit above or below the nearest value existing within acceptable z-score ranges (Tabachnick et al., 2013). For the SM1 cohort, one outlier datapoint was adjusted in each of the GABA+, E/I, go/no-go, and flanker datasets, whilst three outlier datapoints were adjusted in the Stroop data. For the PFC cohort, one outlier datapoint was adjusted in each of the Glx, E/I, and flanker datasets, whilst three outlier datapoints were adjusted in the Stroop data. For aim 1, semi-partial correlations (Spearman’s *rho*) were performed to assess the relationship between age and neurochemical concentrations in the SM1 and PFC regions, residual correcting MRS measures for gender and education to isolate age-effects. For aim 2, partial correlations (Spearman’s *rho*) were performed to assess the relationships between neurochemical concentrations (SM1 and PFC) and inhibitory performance, residual correcting MRS and behavioural measures for age, gender and education to control for demographic-effects. For the exploratory aim, we repeated the semi-partial and partial correlations, as previously described, using the E/I ratio (SM1 and PFC) in place of neurochemical concentrations. To correct for multiple comparisons in the primary analyses (aim 1 and 2), we applied a Bonferroni correction resulting in a corrected alpha for age-neurochemical semi-partial correlations of *p*≤0.025 (2 tests per VOI), and neurochemical-behavioural partial correlations of *p*≤0.013 (4 tests per behavioural measure).

## 3.0 Results

### 3.1 Relationships between healthy ageing and neurochemistry

#### Primary results

As shown in Table 2, after residual correcting for gender and education, no significant correlations were observed between age and GABA+/tCr or Glx/tCr in either the SM1 or PFC regions. To confirm that these null age relationships were not obscured by the residual correction methodology, we performed bivariate spearman’s correlations, which also revealed non-significant results for all comparisons (see Supplemental Material; Table S1).

**Table 2:** Semi-partial correlations between age and neurochemical concentrations.

|  | Age (years) |  |
| --- | --- | --- |
| | Coefficient ($\rho$) | $p$-value (uncorrected) |
| **SM1 Region** |  |  |
| GABA+/tCr | -0.002 | 0.986 |
| Glx/tCr | -0.114 | 0.342 |
| **PFC Region** |  |  |
| GABA+/tCr | -0.021 | 0.891 |
| Glx/tCr | -0.223 | 0.095 |

Neurochemical concentrations were residual corrected for gender and education.

#### Exploratory results

As shown in Table 3, after residual correcting for gender and education, no significant correlations were observed between age and E/I ratio in either the SM1 or PFC regions.

**Table 3:** Semi-partial correlations between age and E/I ratio.

|  | Age (years) |  |
| --- | --- | --- |
| | Coefficient ($\rho$) | $p$-value (uncorrected) |
| **SM1 Region** |  |  |
| E/I ratio | -0.074 | 0.548 |
| **PFC Region** |  |  |
| E/I ratio | -0.116 | 0.440 |

Excitation/inhibition (E/I) ratio was residual corrected for gender and education.

### 3.2 Relationships between neurochemistry and behavioural inhibition

#### SM1 results

Following multiple comparisons correction, results from partial correlations, residual correcting SM1 neurochemical and behavioural measures for age, gender and education, revealed a significant negative relationship between SM1 Glx/tCr concentrations and go/no-go error rates, such that greater concentrations of Glx/tCr in the SM1 region were associated with greater accuracy on the go/no-go (corrected p = 0.034), see Table 4 and Figure 3. SM1 Glx/tCr concentrations were not significantly associated with flanker or Stroop performance (Table 4). After correcting for age, gender, education and multiple comparisons, SM1 GABA+/tCr concentrations were not significantly associated with flanker, Stroop or go/no-go performance (Table 4).

**Table 4:**
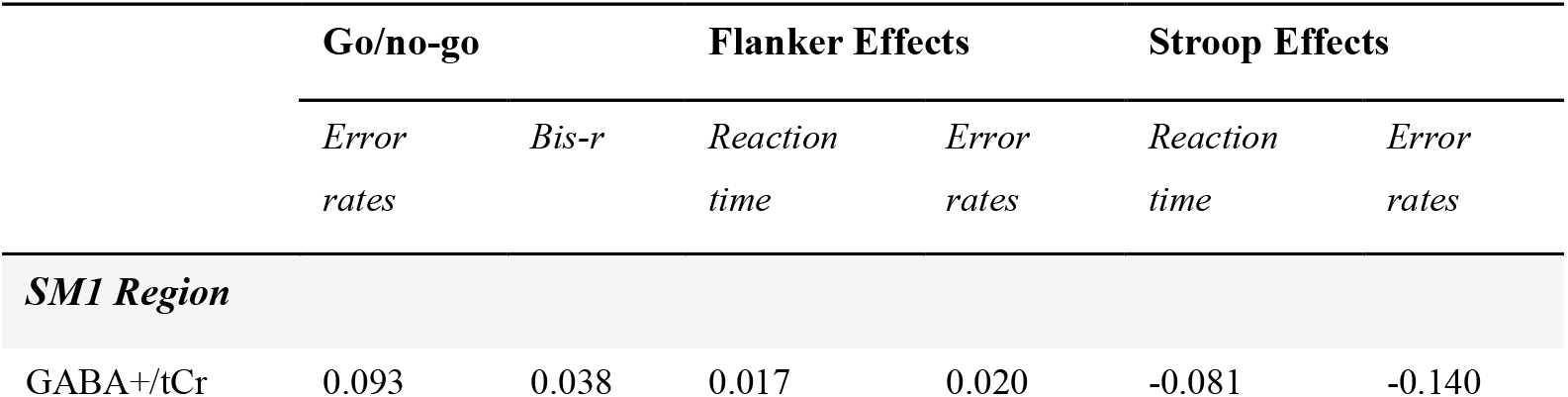
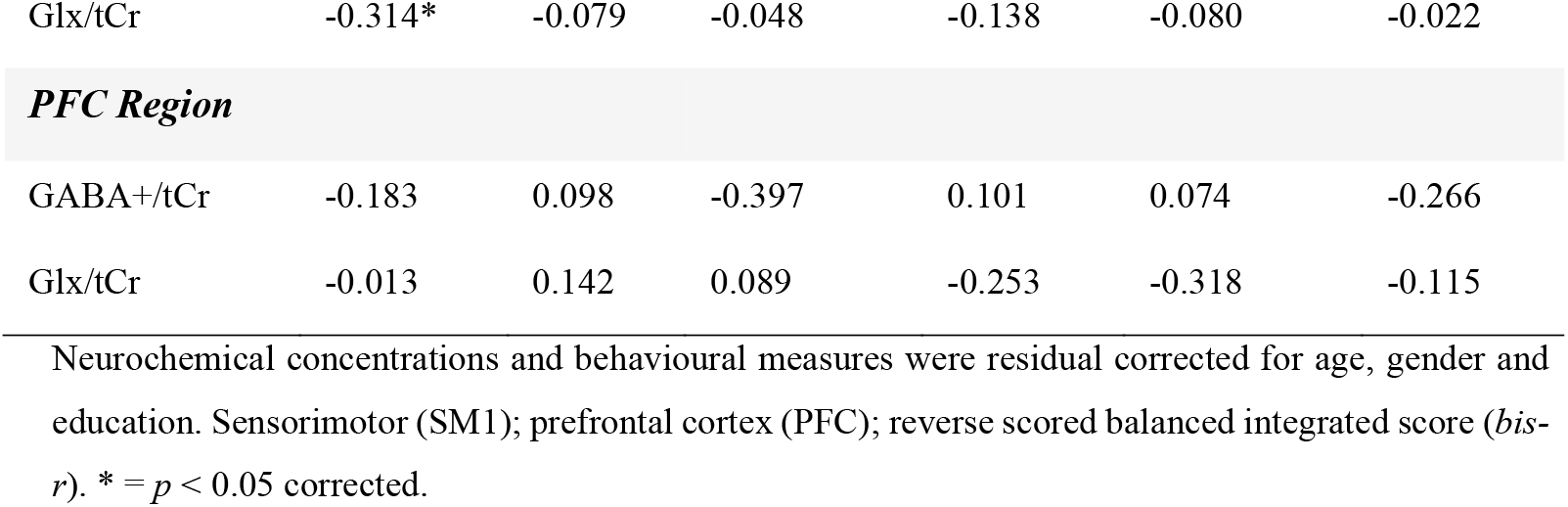
Partial correlations between neurochemical concentrations and behavioural inhibition.

|  | Go/no-go |  | Flanker Effects |  | Stroop Effects |  |
| --- | --- | --- | --- | --- | --- | --- |
|  | Error rates | Bis-r | Reaction time | Error rates | Reaction time | Error rates |
| **SM1 Region** |  |  |  |  |  |  |
| GABA+/tCr | 0.093 | 0.038 | 0.017 | 0.020 | -0.081 | -0.140 |
| Glx/tCr | -0.314* | -0.079 | -0.048 | -0.138 | -0.080 | -0.022 |
| **PFC Region** |  |  |  |  |  |  |
| GABA+/tCr | -0.183 | 0.098 | -0.397 | 0.101 | 0.074 | -0.266 |
| Glx/tCr | -0.013 | 0.142 | 0.089 | -0.253 | -0.318 | -0.115 |

Neurochemical concentrations and behavioural measures were residual corrected for age, gender and education. Sensorimotor (SM1); prefrontal cortex (PFC); reverse scored balanced integrated score (*bis-* *r*). \* = $p < 0.05$ corrected.

**Figure 3:**
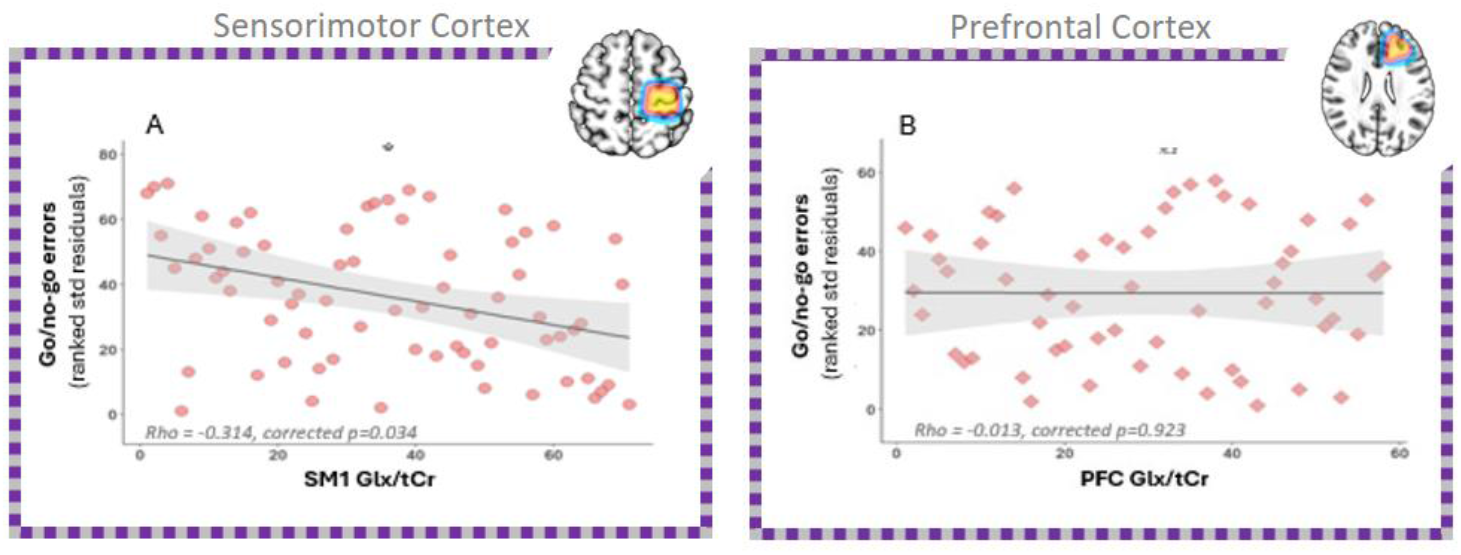
On the left, the primary significant neurochemical-behavioural relationship between SM1 Glx and go/no-go errors (A). On the right, corresponding non-significant relationship in the PFC region (B). SM1 data represented by circular points, PFC data depicted using rhombus points. * = *p* < 0.05 corrected; non-significant (n.s).

#### PFC results

After correcting for multiple comparisons, age, gender and education, no significant partial correlations were identified between PFC GABA+/tCr or Glx/tCr and inhibitory performance, as indexed using the flanker, Stroop and go/no-go (Table 4).

#### Exploratory E/I results

As shown in Table 5, after correcting for age, gender and education, results from partial correlations revealed a significant negative relationship between SM1 E/I ratio and go/no-go error rates, such that a greater E/I imbalance in the SM1 region was associated with greater accuracy on the go/no-go (uncorrected *p* = 0.023). Furthermore, a significant positive correlation was identified between PFC E/I ratio and flanker effects (reaction time), such that a greater E/I imbalance in the PFC region was associated with more pronounced flanker effects (uncorrected *p* = 0.034), see Table 5.

**Table 5:** Partial correlations between E/I ratio and behavioural inhibition.

| Go/no-go |  | Flanker Effects |  | Stroop Effects |  |
| --- | --- | --- | --- | --- | --- |
| Error rates | *Bis-r* | Reaction time | Error rates | Reaction time | Error rates |

| ***SM1 Region*** |  |  |  |  |  |  |
| --- | --- | --- | --- | --- | --- | --- |
| E/I ratio | -0.2777* | -0.0478 | -0.1363 | -0.1725 | 0.0804 | 0.0921 |
| ***PFC Region*** |  |  |  |  |  |  |
| E/I ratio | 0.0984 | -0.0625 | 0.3134* | -0.2213 | -0.2847 | 0.1558 |

E/I ratio and behavioural measures were residual corrected for age, gender and education. Sensorimotor (SM1); prefrontal cortex (PFC); excitation/inhibition (E/I); reverse scored balanced integrated score (*bis-r*). \* = $p < 0.05$ .

## 4.0 Discussion

This MRS study is the first to investigate relationships between sensorimotor and prefrontal GABA+/Glx concentrations and sub-component processes of behavioural inhibition in a healthy ageing population (aged 50-84 years). Surprisingly, we did not identify any relationships between age and GABA+ or Glx concentrations in either the SM1 or PFC regions. This result is incongruent with previous research in healthy older adults, challenging the notion that these neurochemical concentrations invariably decline as a function of age. Importantly, we identified that Glx concentrations in the sensorimotor cortex are significantly associated with a distinct sub-component process of behavioural inhibition, namely response inhibition (as measured using the go/no-go task). For GABA+ from SM1 and PFC, no significant relationships were identified with any of the inhibitory sub-components performance metrics. These MRS findings provide regionally specific evidence that preserving SM1 Glx concentrations benefits response inhibition in healthy older adults, suggesting a role for this neurochemical system in maintaining inhibitory functioning as we age.

### 4.1 No relationship between age and GABA+ or Glx concentrations

In a sample of healthy older adults aged 50-84 years, we did not identify a relationship between age and GABA+ or Glx concentrations in either the SM1 or PFC regions. These results were unexpected, contradicting our first hypothesis. Prior MRS studies have reported an age-related decline in both GABA+ and Glx concentrations across the brain (Porges et al., 2021; Roalf et al., 2020), including motor (Hermans et al., 2018; Kaiser et al., 2005) and frontal (Porges, Woods, Edden, et al., 2017; Rmus et al., 2023) regions. It is important to point out that few MRS-ageing studies have been conducted in healthy older adults (aged ≥50 years). Of these limited studies includes cross-sectional work conducted by Porges and colleagues (2017) who assessed 94 healthy older adults (mean age = 73.1 (±9.9) years) and reported reduced GABA+ concentrations (CSF-correction) in the frontal and posterior (i.e., superior to the splenium and aligned with corpus callosum) brain regions as a function of age. Interestingly, a separate analysis of this data processed with a different tissue correction method did not identify a significant relationship between age and frontal GABA+ concentration in the healthy older adult sample (Porges, Woods, Lamb, et al., 2017). Although various MRS correction method possess certain advantages and disadvantages, with no gold standard, this result highlights that the chosen strategy can impact and/or complicate the interpretation of MRS findings, potentially masking or bolstering age relationships. Furthermore, a recent longitudinal study conducted by Zuppichini et al. (2024) in 30 healthy older adults identified that as age increases SM1/ventrovisual GABA+ concentrations decrease. Interestingly, no significant cross-sectional relationships were identified between age and GABA+ in this cohort, suggesting that assessing within-person age-related neurochemical changes may be more sensitive than group-wide analyses. This reveals another methodological consideration which may provide some explanation for our null age-neurochemical results. Moreover, conflicting results have been reported in younger populations, for example, Mikkelsen et al. (2017) and Aufhaus et al. (2013) did not identify significant age-related changes in GABA+ concentrations in healthy adults aged 18-48 years and 21-53 years, respectively. Altogether, these results highlight that the influence of age on GABA+ is not clear-cut. Additionally, several studies assessing age-related GABA+/Glx changes do so by statistically comparing separate groups of younger and older adults (Grachev & Apkarian, 2001; Hermans et al., 2018; Kaiser et al., 2005; Liu et al., 2024; Rmus et al., 2023; Simmonite et al., 2019). Although this categorical approach elucidates relative, between-group differences, these comparisons do little to characterise ageing trajectories. Indeed, some prior MRS studies have reported age-related declines in GABA+ (age range = 20-76 years; Gao et al. (2013)) and Glx (age-range = 20-70 years; Ding et al. (2016)) with continuity across the lifespan, but a notable gap persists in our understanding of the neurochemical changes occurring in the older adult brain. By placing our results in the context of the existing literature, the seemingly inconsistent ageing results reported here may be influenced by our cross-sectional assessment of healthy older adults specifically (aged ≥ 50 years) and chosen metabolite normalisation method (i.e., referencing to tCr). It’s crucial to reiterate that evidence delineating MRS-assessed neurochemical trajectories in medically/cognitively healthy older adults remains particularly limited. As such, additional cross-sectional and longitudinal multi-regional MRS research in older adult populations, as opposed to between-group assessments, is required to test the prevailing hypothesis of an age-related decline in GABA+ and Glx concentrations. Our null results cast uncertainty on this hypothesis, suggesting that the relationship between older age and neurochemistry may be more nuanced than previously hypothesised.

### 4.2 Glx, but not GABA+, relates with inhibitory sub-components

In the SM1 region, we identified a negative correlation between Glx and go/no-go performance (error rates), indicating that higher SM1 Glx concentrations benefit the ability to inhibit prepotent responding in healthy older adults. This finding extends on recent work by Liu et al. (2024) who reported that higher Glu concentrations correlated with better inhibitory performance (as measured using the SST) in younger adults (n=18; age range = 20-35 years). This MRS result also complements previous fMRI work that demonstrates the SM1 region’s involvement in the suppression of dominant (motor) responding (Aron, 2011), and now suggests a specific role for SM1 neurochemistry in supporting sub-component processes of inhibition during healthy ageing. In contrast, no SM1 Glx relationships were identified with flanker or Stroop performance, which reiterates the importance of considering the non-unitary nature of inhibition when investigating “inhibition” in ageing, as overlooking inhibitory sub-components may inadvertently obscure neurochemical relationships. Overall, response inhibition appears to rely on the neurochemistry of the sensorimotor cortex, having distinct Glutamatergic dependencies. Whilst this is a possibility, it is important to acknowledge that higher Glx concentrations may also be indicative of a healthier neurochemical system more generally speaking. As such, this significant Glx-response inhibition finding could reflect differences in neuronal integrity, whereby individuals with higher Glx relative to group may be due to differences in atrophy and/or tissue composition in the cell. To clarify this alternate explanation, future studies should examine the impact of tissue differences in the investigated VOI’s to better understand the mechanisms driving this Glx association with response inhibition. With respect to SM1 GABA+ concentrations, we did not identify significant relationships with behavioural inhibition measures across tasks. Previous work by Hermans et al (2018) indicates a role for pre-SMA GABA+ in inhibitory functioning in older adults (n=29, mean age = 67.5 (±3.9) years) using an SST paradigm, however, although the SST and go/no-go are often assumed to measure the same form of inhibition (response inhibition), these inhibition tasks seem to rely on distinct neural mechanisms (Raud et al., 2020) which doesn’t support their interchangeable use. This behavioural difference may explain why we did not identify a relationship between GABA+ and behavioural inhibition, in conjunction with our different motor ROI and two-fold larger sample size. Taken altogether, these SM1 results suggest that Glx may play an important role in behavioural inhibition. Although there appears to be a mechanistic focus on GABA+ in ageing literature (Li et al., 2022), our results highlight a relationship between Glx and sub-component processes of inhibition despite the absence of a clear mechanism for this neurochemical system in age-related cognitive decline (Porges et al., 2021). Nonetheless, these neurochemically-and-spatially specific results suggest that preserving Glx in the sensorimotor cortex supports the adequate suppression of inappropriate actions, thereby fostering goal-oriented behaviour crucial for cognition and functional independence. This novel finding suggests that inhibitory (motor) sub-components are sensitive to cerebral SM1 Glx concentration, thus, MRS-assessed Glx may be a useful marker of declining inhibitory functioning during healthy ageing. The neurochemical specificity of these findings lends support to the notion that SM1 Glx is indeed playing a functional role, that said, additional longitudinal work is needed to clarify the role of neuronal integrity and fully elucidate a causal mechanism in the context of healthy ageing.

In the PFC region, before applying Bonferroni corrections for multiple comparisons, we identified weak negative correlations between i) GABA+ and flanker effects (reaction time), and ii) Glx concentrations and Stroop effects (reaction time). These results do underscore that these PFC relationships were not as strong as those identified with Glx in the SM1. Notably, our PFC sample size was considerably smaller than the SM1 because of additional QC and model-fit exclusions (31% reduced for GABA+, 20% reduced for Glx), thus, our PFC analyses were less powered than the SM1 which may explain why these PFC relationships did not surpass corrected statistical thresholds. Altogether, these findings attest that SM1 Glx serves a spatially distinct role in the maintenance of inhibitory functioning, underlining the necessity of considering multiple brain regions to enhance our depth of understanding.

### 4.3 The E/I (im)balance: contradictory evidence

In the present work, utilising the E/I ratio as a measure of neurochemical balance produced differential results. Firstly, we did not detect an ageing influence on the E/I (im)balance. Furthermore, in the SM1 analyses we identified a negative correlation between E/I balance and go/no-go performance (error rates), suggesting that a greater E/I imbalance was associated with greater response inhibition. On the contrary, the PFC E/I balance was positively correlated with flanker performance (reaction time), such that a greater E/I imbalance was associated with more pronounced flanker effects (i.e., poorer inhibition of distractors). These findings effectively contradict one-another, with the SM1 results suggesting that a greater E/I imbalance benefits inhibition whilst the PFC results indicate greater E/I imbalances have a detrimental impact on one’s inhibitory performance. Understanding what the directionality of the E/I (im)balance indicates is subject to interpretation, as higher E/I scores can occur from decreasing GABA+ concentrations and/or increases in the concentration of Glx, or the scores can reflect GABA+ and Glx declining or increasing together at different rates. The GABAergic system has been proposed as primary driver of the E/I imbalance (Bi et al., 2020). We identified the unexpected E/I results in the SM1 (greater E/I imbalance correlates with better go/no-go performance) where we also detected the strong relationship detected between Glx and go/no-go accuracy. Therefore, if GABA+ was indeed driving the E/I imbalance, this may have obscured the SM1 associations since GABA+ did not relate with behavioural inhibition in this region, perhaps potentiating the unexpected directionality of this relationship. Although this is plausible, elucidating the exact source of these contradictory reports is challenging as we are unable to conclusively arbitrate between the possibilities driving E/I imbalances. Overall, the E/I ratio itself represents quite a simplistic measure. The contradictory evidence presented here underscores this ambiguity, which warrants consideration in future research, whilst also highlighting the importance of reporting on the relationships with the neurochemical constituents of the E/I ratio to ensure well-informed conclusions are drawn.

### 4.4 Strengths and limitations

By quantifying GABA+ and Glx concentrations in two separate brain regions (SM1 and PFC) and targeting three inhibitory sub-components, we were able to demonstrate spatially specific relationships between neurochemistry and inhibitory functioning in the context of healthy ageing. However, we do acknowledge several potential limitations. This study was cross-sectional in design which restricts the ability investigate temporal relationships and intraindividual neurochemical changes. The included sample were also particularly healthy, highly functioning and educated, which might not be truly representative of the general older adult population. Although the VOI locations of SM1 and PFC were specifically chosen with respect to the inhibitory tasks, quantifying neurochemical concentrations in other VOIs could have further clarify the spatial specificity of these results. Additionally, our questions were restricted to the study of GABA+, Glx and E/I (im)balance, thus, testing these relationships with other neurochemicals would denote the generalizability of these findings and further characterise healthy brain ageing from a neurochemical perspective. Lastly, we excluded a considerable percentage of the PFC MRS data based on poor data quality (PFC exclusions: 39% of GABA+ and 26% of Glx data). With respect to the VOI placement in left PFC using a large voxel size (3cm^3^), poor data quality was primarily driven by lipid contamination and poor signal/shimming in this frontal region. Although 3cm^3^ voxel dimensions are advised for the HERMES sequence implemented here, we acknowledge that both the large voxel size and the left PFC placement as potential limitations of the current study.

## 5.0 Conclusion

In the present study, we clarified relationships between SM1 and PFC GABA+ and Glx concentrations (as well as their relative balance) and sub-component processes of behavioural inhibition in healthy older adults aged 50-84 years. Contrary to our hypothesis, age was not associated GABA+ or Glx concentrations in either the SM1 or PFC regions, nor did we identify an ageing influence on the relative E/I (im)balance. This unexpected null result casts uncertainty on the prevailing assumption that GABA+ and Glx concentrations decline during healthy ageing. Furthermore, we identified that preserving SM1 Glx concentrations may service an individuals’ capacity to inhibit prepotent responding, suggesting that response inhibition may rely on the Glutamatergic neurochemistry of the sensorimotor cortex. Our utility of the E/I ratio as a measure of neurochemical balance produced differential evidence, with the results from one VOI effectively contradicting the other. Since arbitrating between whether GABA+ or Glx is driving E/I imbalances remains largely inconclusive, utilising this relatively crude measure of neurochemistry warrants careful consideration in future MRS work. Altogether, the present findings suggest a specific role for SM1 neurochemistry in supporting distinct sub-component processes of inhibition, whereby MRS-assessed Glx concentrations may serve as an important indicator of response inhibition over the normal course of ageing.

## 6.0 Data and Code Availability

The study design, hypotheses, and analysis plan were not preregistered. All data pre-processing and analyses were performed using R Statistical Software v4.3.2 (R Core Team, 2023). Data analysis/experiment code and materials are openly available at the project’s Open Science Framework page (osf.io/pbr9f). Data sharing has not yet been approved by the UniSC Human Research Ethics Committee, but the data can be made available upon reasonable request by contacting the corresponding author.

## 7.0 Author Contribution (CRediT)

**Ciara Treacy:** Conceptualisation, Methodology, Formal Analysis, Investigation, Data Curation, Writing-Original Draft, Visualisation, Software, Project Administration. **Sophie C. Andrews:** Conceptualisation, Methodology, Formal Analysis, Writing – Review & Editing, Supervision. **Jacob M. Levenstein:** Conceptualization, Methodology, Software, Formal Analysis, Investigation, Data Curation, Writing – Review and Editing, Visualizations, Supervision, Project Administration, Funding Acquisition

## 8.0 Declaration of Competing Interests

The authors have no competing interests to disclose.

## 9.0 Acknowledgements

This work was completed with support from a Research Training Program Scholarship and SPARK grant (090028321) from the University of the Sunshine Coast. We extend our sincere thanks to the participants who volunteered their time to contribute to this research.

## Notes

https://osf.io/pbr9f/

### Competing Interest Statement

The authors have declared no competing interest.

