## Supplemental Material for "Relationships between GABA+ and Glx concentrations with age and inhibition in healthy older adults"

|  | **Age (years)** | |
| --- | --- | --- |
|  | *Coefficient (rho)* | *p-value (uncorrected)* |
| ***SM1 Region*** |  |  |
| GABA+/tCr | -0.118 | 0.337 |
| Glx/tCr | -0.108 | 0.369 |
| ***PFC Region*** |  |  |
| GABA+/tCr | -0.020 | 0.892 |
| Glx/tCr | -0.249 | 0.062 |

**Table S1:** Bivariate relationships between age and neurochemical concentrations

Neurochemical concentrations were residual corrected for gender and education
